# Genoppi: an open-source software for robust and standardized integration of proteomic and genetic data

**DOI:** 10.1101/2020.05.04.076034

**Authors:** Greta Pintacuda, Frederik H. Lassen, Yu-Han H. Hsu, April Kim, Jacqueline M. Martín, Edyta Malolepsza, Justin K. Lim, Nadine Fornelos, Kevin C. Eggan, Kasper Lage

**Author notes:** Correspondence should be addressed to KCE and KL. These authors contributed equally (Greta Pintacuda, Frederik H. Lassen, Yu-Han H. Hsu, April Kim).

## Abstract

Combining genetic and cell-type-specific proteomic datasets can lead to new biological insights and therapeutic hypotheses, but a technical and statistical framework for such analyses is lacking. Here, we present an open-source computational tool called Genoppi that enables robust, standardized, and intuitive integration of quantitative proteomic results with genetic data. We used Genoppi to analyze sixteen cell-type-specific protein interaction datasets of four proteins (TDP-43, MDM2, PTEN, and BCL2) involved in cancer and neurological disease. Through systematic quality control of the data and integration with published protein interactions, we show a general pattern of both cell-type-independent and cell-type-specific interactions across three cancer and one human iPSC-derived neuronal type. Furthermore, through the integration of proteomic and genetic datasets in Genoppi, our results suggest that the neuron-specific interactions of these proteins are mediating their genetic involvement in neurodevelopmental and neurodegenerative diseases. Importantly, our analyses indicate that human iPSC-derived neurons are a relevant model system for studying the involvement of TDP-43 and BCL2 in amyotrophic lateral sclerosis.

## Main Text

Large genetic datasets, such as those obtained from genome-wide association studies (GWAS) or exome sequencing are becoming increasingly available. Simultaneously, advanced proteomic technologies can generate high-quality cell- and tissue-specific quantitative proteomic data (e.g., from immunoprecipitations followed by tandem mass spectrometry [IP-MS/MS]^1,2^ or whole-proteome analyses of cells or tissues^3^). Integration of genomic and proteomic datasets has revealed that genetic variation implicated in rare and common diseases often manifests at the proteome level, for example, by impacting protein complexes or cellular networks^4^. However, while data from genetics and cell-type-specific quantitative proteomics have the potential to inform each other and lead to key molecular and biological insights, the relevant data types are often not interoperable, even for experts in these fields. This creates a bottleneck for interpreting genetic data and learning about cell-type-specific molecular biology of human diseases. In the longer term, difficulties in reconciling these data types hamper efforts towards gaining mechanistic insights from genetic data and designing therapeutic interventions. To enable the robust, standardized, and intuitive integration of data from cell-type-specific quantitative proteomics and genetics, we have developed an open-source computational tool named Genoppi (github.com/lagelab/Genoppi).

Given log_2_ fold change (FC) values between studied conditions (e.g., bait vs. control IPs) for multiple experimental replicates, Genoppi identifies statistically enriched proteins by applying user-defined threshold, displays the data in scatter and volcano plots, and provides options for integrating the results with other types of data (**Fig. 1a, Online Methods** and **Supplementary Protocol**). Genoppi provides functionality to quality control (QC) a protein interaction dataset^5^ by testing whether it is enriched for known interaction partners compiled from >40,000 scientific articles into the protein interaction network InWeb_InBioMap^6^ (**Fig. 1b**). The ability to automatically integrate InWeb_InBioMap with experimental data in real-time makes it easy to distinguish between published versus newly identified interaction partners of a protein of interest in the context of a new interactome study (without the need to extensively interrogate the literature on a case-by-case basis).

**Figure 1.**
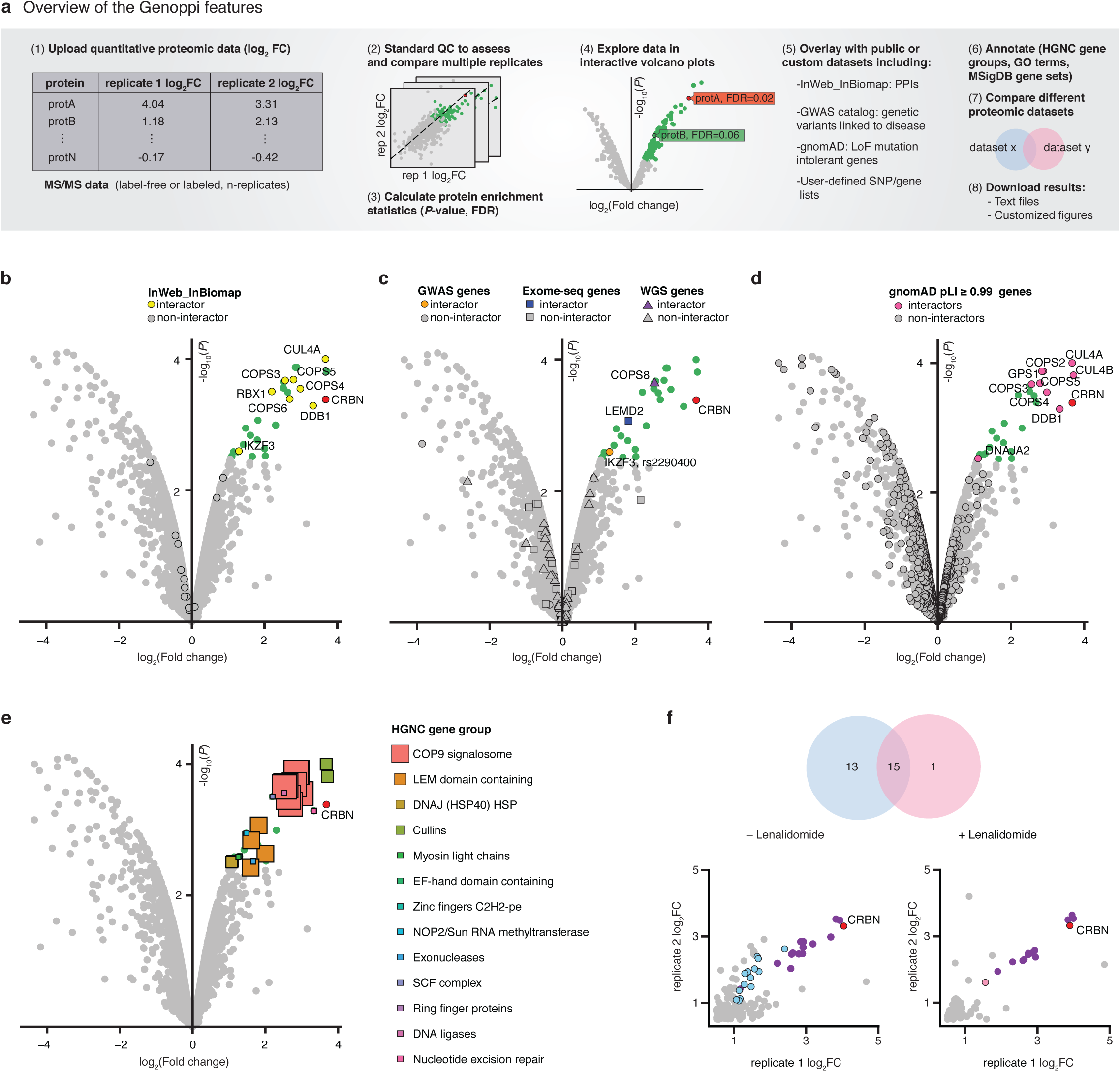
Overview of Genoppi. **(a)** Overview of the Genoppi features. FC, fold change; MS, mass spectrometry; QC, quality control; FDR, false discovery rate; PPI, protein-protein interaction; LoF, loss-of-function; SNP, single nucleotide polymorphism. **(b)** Volcano plot of published CRBN interaction data in MM1S multiple myeloma cells versus control samples. The x-axis shows the log_2_ FC of each identified protein and the y-axis the corresponding -log_10_ *P*-value. The bait protein (CRBN) is marked in red; statistically enriched interactors with log_2_ FC > 0 and FDR ≤ 0.1 are in green; non-interactors that do not pass this threshold are in grey. Known interactors of CRBN in InWeb_InBioMap are marked by black border circles; those enriched in the experimental data are highlighted in yellow (overlap enrichment *P* = 2.1e-12). **(c)** The volcano plot from (b) is overlaid with genetic data. Proteins encoded by genes mapped from acute lymphoblastic leukemia GWAS SNPs (GWAS genes, significantly mutated genes identified through exome sequencing in multiple myeloma (Exome-seq genes), or recurrently mutated cis-regulatory elements identified via whole-genome sequencing in multiple myeloma (WGS genes) are marked by black border circles, squares, or triangles, respectively; those enriched in the experimental data are highlighted in orange, blue, or purple, respectively. **(d)** The volcano plot from (b) is overlaid with proteins intolerant of LoF mutations in gnomAD. Proteins encoded by genes with pLI scores ≥ 0.99 are marked by black border circles; those enriched in the experimental data are highlighted in magenta. **(e)** The volcano plot from (b) is overlaid with HGNC gene group annotations (square markers) for the enriched interactors. Marker size scales with the number of interactors assigned to each group. **(f)** Comparison of CRBN interaction partners in untreated (-Lenalidomide) versus lenalidomide-treated (+Lenalidomide) MM1S cells. Top: Venn diagram representing the overlap of enriched (log_2_ FC > 0 and FDR ≤ 0.1) interactors between the two conditions. Bottom: scatter plots showing log_2_ FC of identified proteins in two replicates (x- and y-axis, respectively) for each condition. Interactors shared between conditions are shown in purple; interactors unique to each condition are labeled in blue or pink, respectively.

Genoppi can also integrate various types of genetic data. One such data type is a list of single nucleotide polymorphisms (SNPs) derived from GWAS of a disease of interest. Mapping SNPs to genes and, subsequently, to specific proteins encoded by these genes, is technically nontrivial. For this reason, Genoppi automatically uses linkage disequilibrium (LD) information from the 1000 Genomes Project^7^ to identify proteins in a proteomic dataset that are encoded by genes in LD loci of NHGRI-EBI GWAS catalog^8^ or user-defined SNPs^9^ (**Fig. 1c** and **Online Methods**). In addition, exome and whole-genome sequencing has been increasingly used to identify genes that have a significant burden of rare mutations in patients with a particular disease compared to healthy individuals. Genoppi is designed to enable the identification of proteins in a proteomic dataset based on user-defined gene lists derived from such studies^10^ (**Fig. 1c**). Genoppi can also incorporate gene constraint data from gnomAD^11^ to label proteins intolerant to loss-of-function mutations (**Fig. 1d**). When these external datasets are integrated with proteomic results, overlaps are displayed as Venn diagrams and statistically tested when appropriate (**Online Methods** and **Supplementary Protocol**).

Another feature of Genoppi is annotation of proteomic data with gene sets from several databases, including HGNC^12^, GO^13,14^, and MSigDB^15,16^, to visually identify groups of proteins that may be overrepresented in a particular dataset (**Fig. 1e**). Finally, it is possible to perform head-to-head comparisons of proteomic experiments performed under different conditions using Genoppi. For example, an interaction experiment executed with and without drug treatment^5^ (**Fig. 1f**), or between the wild-type and mutated protein version of an oncogene, to elucidate the cellular effects of either pharmaceutical or genetic perturbations in a particular cell type. Overall, Genoppi provides various ways to explore proteomic datasets, to guide hypothesis generation, or to inform targeted and cost-efficient follow-up experiments.

Genoppi is an open-source software that is easily accessible and flexible to custom needs in the research community. In particular, the Genoppi web application provides a simple interactive interface with customizable options and the ability to work with a wide variety of different types of quantitative proteomic datasets (e.g., IP-MS/MS analyses, or whole-proteome analyses of cells or tissues). For example, it is possible to dynamically explore a dataset by changing various technical thresholds and getting results in real-time; furthermore, a search function enables quick identification of proteins of interest in various plots (**Supplementary Protocol**). Users can locally download the generated data and plots to share with collaborators. For users interested in building custom analytical pipelines, Genoppi functions are also available as an R package with detailed documentation. In summary, Genoppi can considerably facilitate data sharing and analysis in cross-disciplinary collaborations that are now common in both academia and industry. Further details about options, workflows, and analyses can be found in **Supplementary Protocol**.

To exemplify how analyzing and comparing cell-type-specific proteomic data using Genoppi can uncover convergent and divergent disease-relevant biology of the same protein in different cell types, we generated IP-MS/MS data for four proteins of interest (TDP-43, PTEN, BCL2, MDM2; baits, hereafter) in four distinct cell lines (**Online Methods** and **Supplementary Table 1**). We chose these four baits because they play important, but not fully elucidated roles in cancer, neurological disease and psychiatric conditions.

We executed IPs in a human iPSC-derived neuronal cell line (glutamatergic patterned induced neurons [GPiNs]^17^) and three cancer cell lines (G401, T47D, and A375; **Fig. 2a-c, Supplementary Fig. 1a**), along with control experiments using isotype-matched immunoglobulin gamma (IgG; **Online Methods**). All IP and IgG control experiments were conducted in triplicate and quantitated by liquid chromatography followed by label-free tandem mass spectrometry (LC-MS/MS; **Online Methods**). Genoppi was used to: i) quality control, plot, and visualize the IP-MS/MS results; ii) identify enriched interaction partners of each bait in each cell type; iii) compare enriched interaction partners between cell types; and iv) integrate these data with genetics for biological discovery and hypothesis generation (**Fig. 2, Supplementary Fig. 1, Supplementary Table 2**, and **Online Methods**).

**Figure 2.**
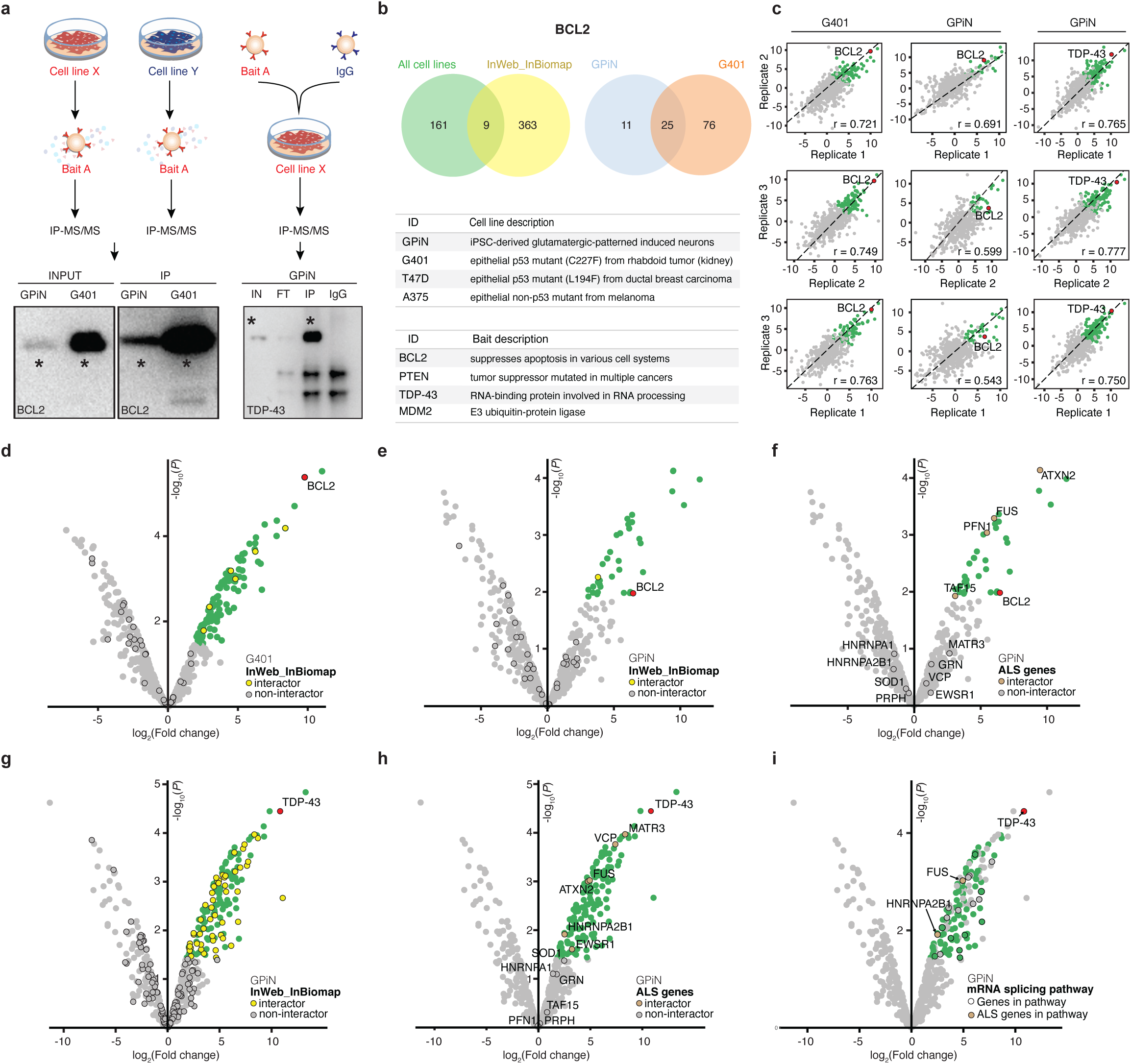
IP-MS/MS analysis through Genoppi. **(a)** Experimental design and representative western blots of immunoprecipitations for MS/MS experiments; a star indicates the molecular weight (MW) of the bait (BCL2 and TDP-43, respectively); IN, FT, IP, and IgG stand for input, flow-through, immunoprecipitation, and IgG isotype control, respectively. **(b)** Top: Venn diagrams representing the overlap between BCL2 interactors identified in all cell lines and known InWeb_InBioMap interactors, and the overlap of interactors identified in neurons (GPiN) and a cancer cell line (G401). Bottom: complete list of cell lines and baits used for the experiments. **(c)** Scatter plots showing reproducibility of three IP replicates in terms of log_2_ FC correlation for three sets of experiments: BCL2 versus IgG control in G401 cells or GPiNs, and TDP-43 versus IgG control in GPiNs. Pearson correlation (r) is reported in each plot. **(d)** BCL2 versus IgG control IP results in G401 cells. The volcano plot is overlaid with known BCL2 interactors in InWeb_InBioMap (overlap enrichment *P* = 0.61). **(e)-(f)** BCL2 versus IgG control IP results in GPiNs. The volcano plot is overlaid with known BCL2 interactors in InWeb_InBioMap (e; overlap enrichment *P* = 0.91) or proteins encoded by ALS genes (f; overlap enrichment *P* = 0.005). **(g)-(h)** TDP-43 versus IgG control IP results in GPiNs. The volcano plot is overlaid with known TDP-43 interactors in InWeb_InBioMap (g; overlap enrichment *P* = 0.04) or proteins encoded by ALS genes (h; overlap enrichment *P* = 0.07). In plots (c)-(h), the bait (BCL2 or TDP-43), interactors (log_2_ FC > 0 and FDR ≤ 0.1), and non-interactors are shown in red, green, and grey, respectively; overlaid proteins are marked by black border circles. **(i)** TDP-43 versus IgG control IP results in GPiNs shown as volcano plot, with interactors (log_2_ FC > 0 and FDR ≤ 0.1) that are GPiN-specific (i.e. not interactors in G401) shown in green; other detected proteins are in grey; bait (TDP-43) is in red. Black border circle indicates interactors in the MSigDB Reactome mRNA splicing pathway; two GPiN-specific interactors in the pathway, FUS and HNRNPA2B1, have been linked to ALS and are highlighted in brown.

We first explored the analysis results for BCL2, which is a well-studied oncogene functioning as an apoptosis suppressor in a variety of cell types^18^; it was also recently shown to be an important regulator of plasticity and cellular resilience during neuronal development^19^. Intriguingly, data from both human brain tissue and animal models of neuropathological conditions suggest a role for BCL2 in cell death regulation in the mature nervous system^20^ and in amyotrophic lateral sclerosis (ALS)^21,22^. However, little is known about the specific role of BCL2 in different human cell types, and insights into its overlapping and differential interaction partners in neurons compared to cancer cell lines could generate actionable biological hypotheses of therapeutic relevance.

Across the four tested cell lines, BCL2 had a total of 170 nonredundant statistically enriched interaction partners. Interaction partners of any bait protein will hereafter be defined as proteins with a log_2_ FC > 0 and false discovery rate (FDR) ≤ 0.1 in the experiment compared to the cell-type-matched IgG control (**Online Methods**), while non-interactors are proteins that do not fulfil either or both criteria. Amongst BCL2 interactors only nine were previously reported in InWeb_InBioMap (**Fig. 2b** and **Supplementary Table 2**). However, when comparing the BCL2 interaction partners in GPiNs versus a cancer cell line (G401), we found both a large set of shared interactors and interactors unique to each cell line (**Fig. 2b**). Specifically, 25 out of 36 (69%) interactors found in GPiNs were also interactors in G401 cells; conversely, 25 out of 101 (25%) interactors found in G401 cells were also interactors in GPiNs (*P* = 2.6e-12, using a hypergeometric distribution). This supports the observation that, despite little overlap with InWeb_InBioMap interactors and the identification of many cell-type-specific interactors (11 in GPiNs and 76 in G401 cells, respectively), a significant subset of new BCL2 interaction partners (n = 24) were common to two different cellular backgrounds and are likely true biological interactors (**Supplementary Table 2**).

We next determined the correlation between replicate samples (**Fig. 2c**) and visualized the differential interactions of BCL2 in G401 cells versus GPiNs (**Fig. 2d-f**). Superimposing known interaction partners from InWeb_InBioMap with proteins detected in our IPs (**Fig. 2d,e**) shows that the majority of the interaction partners we identified in both cell types are new (35 of 36 in G401 cells and 95 of 101 in GPiNs; **Supplementary Table 2**).

To test whether the BCL2 interactome was enriched for known cancer driver genes in G401 cells and for genes involved in neurological disorders in GPiNs, we integrated genetic data with proteomic data in Genoppi. We used datasets of cancer driver genes^23^, genes involved in neurodevelopmental delay, autism spectrum disorders^24,25^ and schizophrenia^26^, as well as a curated list of genes involved in ALS^27,28^ (**Supplementary Table 3**). The statistical analyses compare the enrichment of disease-related proteins amongst the interactors of the bait protein versus the non-interactors (**Online Methods**). In this way, the statistical enrichment is always conditional on the proteome being expressed in the tested cell line.

We found that known cancer driver genes were evenly distributed between proteins significantly interacting with BCL2 and the rest of the G401 proteome (i.e., the non-interactors; **Supplementary Fig. 2a**). In contrast, the BCL2 interactome in neurons was significantly enriched for ALS-associated proteins compared to the overall GPiN proteome (*P* = 0.005, using a hypergeometric distribution; **Fig. 2f**). We further show that ALS-implicated proteins were not enriched among BCL2 interactors found in any of the three cancer cell lines (**Supplementary Fig. 2b-d**), confirming that a connection to ALS is a feature of the neuron-specific interaction partners of BCL2. Genoppi thus points to targeted follow-up experiments to guide the characterization of newly identified neuron-specific interactors of BCL2.

TDP-43, is a ubiquitously expressed RNA-binding protein encoded by the gene *TARDBP*. Rare mutations in *TARDBP* have been identified as a cause of familial ALS and Frontotemporal dementia (FTD). TDP-43 aggregation is also a common pathological hallmark of both neurodegenerative disorders^29^. To gain insights into neuronal functions of TDP-43, we analyzed the IP-MS/MS data of TDP-43 in GPiNs (**Fig. 2a-c**) and found many of its known, InWeb_InBioMap interaction partners (*P* = 0.04, using a hypergeometric distribution; **Fig. 2g**). We also integrated known ALS-associated risk genes with the TDP-43 proteome in GPiNs and show that the experiments in GPiNs recapitulate known interactions between TDP-43 and MATR3, VCP, FUS, ATXN2, HNRNPA2B1, and EWSR1 (**Fig. 2h**).

In the literature, it is highly debated whether the pathologies related to TDP-43 aggregation are due to altered transcriptional regulation or cell toxicity. Interestingly, a large portion of TDP-43 interactors in GPiNs (including four out of six detected ALS-associated proteins) are involved in RNA-metabolism, suggesting that the role of TDP-43 in human neurons is mostly related to transcriptional and post-transcriptional regulation, as highlighted using Genoppi’s gene set annotation feature (**Fig. 2i, Supplementary Protocol**). Importantly, none of the ALS-associated proteins involved in transcriptional or post-transcriptional regulation were identified among TDP-43 interactors in any of the cancer lines (**Supplementary Table 4**), suggesting that in non-neuronal contexts, TDP-43 may function through a different molecular mechanism involving a separate, cell-type-specific set of interactors.

TDP-43 has previously only been studied in human brain homogenates, which are a mixture of many different cell types^30^, or in HEK cells^31^ that are less relevant to its role in ALS or FTD. Our results illustrate that GPiNs recapitulate known biology of this protein and its physical interactions to a number of known ALS-related proteins. Together, our findings support the idea that human iPSC-derived neurons are a suitable model for studying TDP-43 in the context of ALS and provide testable hypotheses about its interaction with proteins involved in neurodegenerative diseases. In the future, it will be of interest to test the functional significance of the convergence of ALS risk genes in the TDP-43 pathway and dissect the role of individual interactions between TDP-43 and ALS-related proteins in the context of transcript regulation.

We performed analogous experiments and Genoppi analyses of two more proteins (MDM2 and PTEN) that are also hypothesized to have divergent functions in cancer and neurodevelopment. Similar to the observations for BCL2 and TDP-43, we observe cell-type-specific interaction patterns that inform targeted hypotheses and follow-up experiments (**Supplementary Note 1, Supplementary Fig. 1b-h**, and **Supplementary Table 2**).

Combining the IP-MS/MS results of BCL2, TDP-43, MDM2 and PTEN across four different cell lines showed that only 15.6% of the interaction partners we identified are reported in InWeb_InBioMap, meaning that up to 84.4% of these interactions are new, offering potentially exciting insights into the biology of these proteins. Stratified per protein, 97.3%, 94.8%, 71.7%, and 64.5% of the interaction partners were new for PTEN, BCL2, MDM2, and TDP-43, respectively. Across cell lines, 84.4%, 83.5%, 82.1%, and 80.9% of the interaction partners identified for the four proteins are new in G401, A375, GPiN, and T47D cells, respectively.

In **Fig. 2b**, we show how newly identified interaction partners of BCL2 in G401 cells can be replicated in GPiNs supporting the biological valididity of these interactions even if they have not been previously reported in the literature. We extended the same analysis to all four baits in all possible pairs of cell lines (**Supplementary Fig. 3**) showing that although we identified a large set of new interaction partners for each bait, they can be reproduced in multiple cell types. In addition, when clustering cell lines based on the pairwise overlap of significant interaction partners for TDP-43, BCL2 and PTEN, we observe a clear clustering of cancer cells (G401, A375, and T47D) versus GPiNs (**Supplementary Fig. 3**). This indicates that, as expected, the cancer cell interactomes are more similar to each other than to the neuronal interactomes for three out of four proteins tested.

Finally, we pooled the significant interaction partners of all four bait proteins in the three cancer cell lines (cancer cell interactors, hereafter) and in the neurons (GPiN interactors, hereafter), respectively. While the cancer cell interactors were not enriched for cancer driver genes, the GPiN interactors were significantly enriched for the ALS genes (*P* = 0.004, using a hypergeometric distribution).

Together, all four tested proteins exhibited a pattern of both unique interaction partners in each cell type and a statistically significant set of shared interaction partners across cell types. Our results further suggest that the neuron-specific interactions of BCL2 and TDP-43 are associated with genetic ALS disease risk and its related biology. Overall, our data indicate that proteins have both cell-type-specific and cell-type-independent interactions and that are both relevant to genetic diseases.

Excellent programs are available to the community to analyze raw tandem mass spectrometry data^32-34^, while other tools, such as ProHits-viz^35^, provide outstanding visualization capabilities to summarize protein interaction data as well as communicate quantitative differences between a protein of interest and its potential interaction partners. However, none of these tools focuses on creating a systematic and unified workflow particularly aimed at integrating cell-type-specific quantitative proteomic datasets and genetic information. Genoppi is designed so it can be easily incorporated in any functional genomics pipeline by allowing users to integrate datasets, download the results, and modify the code as needed to extend the software and meet different usability requirements.

In this paper, we showcase Genoppi as a versatile tool that can be employed to combine original and published datasets in a simple and clear format, allowing easy analysis, visualization and exploration of otherwise heterogeneous proteomic datasets. We believe that as more genetic and proteomic datasets become available, Genoppi will become increasingly valuable.

### Data availability

All processed IP-MS/MS data generated in this study are available in **Supplementary Table 5**.

### Code availability

Source code and documentation for the Genoppi R package and Shiny application are available on GitHub (https://github.com/lagelab/Genoppi). Custom Python and R scripts to pre-process the IP-MS/MS data and generate the figure panels in this study are available from the corresponding authors upon request.

## Acknowledgements

We thank Zuzana Tothova, Josephine Kahn, Siddhartha Jaiswal, and Srinivas Viswanathan for discussions on data analysis and J.T. Neal for providing the cancer lines used in this study. We are grateful to Monica Schenone, Jake Jaffe, Karl Clauser, D.R. Mani and Steve Carr from the Broad Institute Proteomics Platform for explaining proteomic data structures, discussions of analyses, and for providing help interpreting the Spectrum Mill output. This work was supported by grants from The Stanley Center for Psychiatric Research, the National Institute of Mental Health (R01 MH109903), the Simons Foundation Autism Research Initiative (award 515064), the Lundbeck Foundation (R223-2016-721), National Institute of Diabetes and Digestive and Kidney Diseases (T32DK110919), and a Broad Next10 grant.

## Author contributions

GP, KCE, KL designed the experiments. AK, EM, JL developed algorithms with supervision from KL. FHL, YHH, AK developed the Genoppi R package and application. GP and JMM performed the experiments. GP, FHL, YHH, AK, NF, KCE, KL analyzed the experimental data. GP, FHL, YHH, AK, JMM, NF, KCE, KL wrote the paper with input from all authors. KL initiated and led the project.

## Online Methods

A user-friendly documentation of analytical and visualization features implemented in the Genoppi application is provided in **Supplementary Protocol**. This section provides additional technical details for analyses performed by Genoppi.

### Moderated t-test for identifying enriched interactors

Given protein enrichment values, in terms of log_2_ fold change (FC), from ≥ two replicates, Genoppi performs a one-sample moderated *t-*test from the limma^36^ R package to calculate a two-tailed *P*-value and Benjamini-Hochberg false discovery rate (FDR) for each protein enrichment value. Limma was originally developed to robustly identify differentially expressed genes in microarray experiments and has since been used on a variety of data types, including proteomic results^37,38^. The empirical Bayes moderated *t*-test is used in Genoppi, as it is less sensitive to underestimated sample variances and performs best on small sample sizes compared to the classical *t*-test^37^.

### SNP-to-gene mapping

To generate the pre-calculated data Genoppi uses for SNP-to-gene mapping, we filtered the 1000 Genomes Project^7^ Phase 3 dataset to obtain genotype data for unrelated individuals residing in Utah with Northern and Western European ancestry and SNPs with minor allele frequency ≥ 0.05 and missing rate ≤ 0.1. Pairwise linkage disequilibrium (LD) between SNPs was calculated using a sliding window of 200kb, which is the default haplotype block estimation distance used in PLINK^39^. Next, for each SNP, the LD genomic locus was defined as the region covered by other SNPs that have r^2^ > 0.6 with the SNP, ± 50kb on either end. These parameters were chosen to comply with established community standards^26,40-42^. Using the pre-calculated LD locus boundaries, Genoppi can then efficiently identify all Ensembl^43^ protein-coding genes whose coordinates overlap with LD loci given a SNP list of interest. If multiple genes are present in the locus defined by a SNP of interest, all genes are mapped to that SNP.

To verify that SNPs are robustly mapped to genes using the mapping method in Genoppi, we mapped 20 random SNPs to genes using both Genoppi and Disease Association Protein-Protein Link Evaluator (DAPPLE)^44^, which is a standard tool for SNP-to-gene mapping. DAPPLE uses the following definition for LD locus of a SNP: “the region containing SNPs with r^2^> 0.5… extended to the nearest recombination hotspot.” Nonetheless in our comparison test, 100% of genes mapped from SNPs using DAPPLE were analogously mapped using the Genoppi algorithm, illustrating the robustness of our approach.

### Hypergeometric test for assessing overlap between datasets

One-tailed *P*-values are calculated using a hypergeometric distribution to identify the significance of overlap between experimental proteomic results and other genetic datasets, known protein interactors from InWeb_InBioMap^6^ (InWeb), or genes intolerant of loss-of-function (LoF) mutations derived from gnomAD^11^. To test overlap with a genetic dataset (e.g., known causal genes for a disease), the “population” (*N*) is defined as all genes encoded by proteins identified in the experimental data and “success in population” (*k*) is defined as the subset of *N* that passed the user-defined significance threshold (i.e., statistically enriched genes). The “sample” (*n*) contains genes from the genetic dataset that are found in *N* and “success in sample” (*x*) is the overlap between *k* and *n*. Similar definitions apply to test overlap with InWeb interactions or gnomAD genes, except in this case the “population” (*N*) is the intersection of all genes in the experimental data and in the respective database, while the “sample” (*n*) is the subset of *N* consisting of InWeb interactors for a chosen bait or LoF-intolerant genes defined using a gnomAD pLI score cutoff (**Supplementary Protocol**). The hypergeometric test^45^ is performed to calculate the statistical significance of having a given amount of success in a population. This procedure tests the statistical significance of an overlap between the proteomic and external datasets, while taking into consideration that only a subset of all proteins (and their corresponding genes) are identified in the proteomic data, meaning that the statistical test is conditional on the proteome of the cell type being tested.

The hypergeometric test is not performed for genes derived from the SNP-to-gene mapping feature in Genoppi. To correctly test the overlap between these genes and the proteomic results is a complicated statistical problem that can easily lead to confounded results and inflated *P*-values. Confounders include whether the mapped gene is from a single or multigenic locus, the gene length, its tissue-specific expression pattern, to name a few. To accurately perform this analysis requires a workflow that is dataset-specific and is beyond the scope of Genoppi. To not mislead users, and to ensure that other statistical tests in Genoppi can be considered reliable, we do not test the statistical significance of overlap between proteomic data and genes mapped from SNPs.

## Experimental procedures

### Cell cultures

#### Glutamatergic patterned induced neurons (GPiNs)

GPiNs were differentiated from an induced pluripotent stem cell (iPSC) line (human dermal fibroblast, neonatal, HDFn) by conditional expression of the neuralizing transcription factor NGN2^17^. A plate was coated with GelTrex (LifeTechnologies, A1413301) adhesion matrix (1:100 in DMEM/F:12) and cells were seeded at a density of 40,000 cells cm^-2^ in Stemflex media (Gibco, A3349401) containing Genetecin as selective antibiotic (Thermo Fisher, 10131027) (1:400). Once the cells achieved 60% confluency, the monolayer was detached using Accutase (Gibco, A11105) and passaged onto new plates for cell expansion.

#### Cancer cell lines

We used the following cancer cell lines: A375 (ATCC CRL-1619), a human malignant melanoma cell line exhibiting a wild-type p53 genotype; G401 (ATCC CRL-1441), a kidney rhabdoid tumor cell line with a wild-type p53 genotype and T47D (ATCC HTB-133), a human breast tumor cell line with a mutated p53 genotype. All cell lines were plated at a density of 40,000 cells cm^-2^ (uncoated plates). Cell maintenance media contained 10% FBS and PenStrep (1:1000). A375 cells were cultured in DMEM (Thermo Fisher) and split every two days (at a 1:12 ratio). G401 were cultured using McCoy’s 5A (Thermo Fisher) media and split every two days (at a 1:10 ratio), and T47D cells were cultured in RPMI no phenol red (Thermo Fisher) media and split every three days (at a ratio of 1:4). All cell lines were incubated at 37°C, 5% CO_2_. To achieve detachment during passaging, all cell lines were exposed to TrypLE (Thermo Fisher).

### Differentiation of GPiNs

HDFn cells, stably expressing TetO-Ngn2-Neo and reverse tetracycline-controlled transactivator (rtTA), were plated at a density of 40,000 cells cm^-2^ with rock inhibitor Y27632 (Stemgent, 04-0012). Day 1 cells were differentiated in N2 media (Gibco) supplemented with 10 μM SB431542 (Tocris, 1614), 2 μM XAV939 (Stemgent, 04-00046) and 100 nM LDN-193189 (Stemgent, 04-0074) along with doxycycline hyclate (2 μg mL^-1^). Day 2 media was N2+SB/XAV/LDN/doxycycline hyclate and differentiation media were as previously described^46^. On day 3, cell differentiation was continued in neurobasal media (Gibco) supplemented with B27 (50X, Gibco), brain-derived neurotrophic factor (BDNF), ciliary neurotrophic factor (CTNF), glial cell-derived neurotrophic factor (GDNF) (R&D Systems 248-BD/CF, 257-NT/CF, and 212-GD/CF at 10 ng mL^-1^) and doxycycline hyclate (2 μg mL^-1^).

### Protein extraction and immunoblotting

Total protein extract was obtained by harvesting cells and either processing them immediately or snap-freezing them on dry ice for storage at -80°C^47^. In both cases, cell pellets were washed with PBS and resuspended in 10x packed cell volume (PCV) IP lysis buffer (Pierce), with freshly added Halt protease and phosphatase inhibitors (Thermo Fisher). After a 20 min incubation time at 4°C, cells were collected by centrifugation (16,200 g, 20 min, 4°C) and resuspended in 3x PCV lysis buffer. The concentration of the samples was quantified using the Thermo BCA protein assay and when not used immediately, samples were stored at -80°C. Samples for immunoblotting were diluted in 6xSMASH buffer (50 mM Tris HCl pH 6.8, 10% glycerol, 2% SDS, 0.02% bromophenol blue, 1% b-mercaptoethanol), boiled for 10 min at 95°C, separated on a NuPAGE 4-12% Bis-Tris Protein Gel (Thermo Fisher), and transferred onto a PVD membrane (Thermo Fisher) by wet transfer (100 V for 2 hours). Membranes were blocked by incubation for one hour at room temperature in 10 mL TBS and 0.1% Tween (TBST) with 5% w/v BioRad Blotting-grade Blocker. Blots were incubated overnight at 4°C with the primary antibody, washed 3 times for 10 min with TBST and incubated for 45 min with secondary antibody conjugated to horseradish peroxidase. After washing 3 times for 5 min with TBST, bands were visualized using ECL (GE Healthcare). All antibodies used in this study are listed in **Supplementary Table 1.**

### Immunoprecipitations

For each individual experiment, 1-2 mg of protein extract was incubated at 4°C overnight in the presence of 1-2 μg of the relevant antibody. On the next day, 50 μL of Protein G beads (Pierce) were added to each sample and incubated at 4°C for 4 hours. Flow-through was collected and beads were washed once with 1 mL lysis buffer (Pierce) supplemented with Halt protease and phosphatase inhibitors (Thermo Fisher), and twice with PBS. Beads were resuspended in 60 μL of PBS and 10% of the volume was employed for immunoblotting, after being boiled in 6xSMASH buffer (50 mM Tris HCl pH 6.8, 10% Glycerol, 2% SDS, 0.02% bromophenol blue, 1% b-mercaptoethanol) for 10 min at 95°C. The remaining volume was stored at -80°C and subsequently used for mass spectrometry (MS) analysis.

### Sample preparation for mass spectrometry

All immunoprecipitated samples (*n*=48) and IgG controls (*n*=12) were in PBS buffer on beads. PBS was removed and samples were dissolved in 50 µL TEAB (tri ethyl ammonia bicarbonate, 50 mM) buffer, followed by trypsin (Promega) digestion for 3 hours at 38°C. Digested samples were dried to 20 µL and 10 µL were subjected to MS analysis.

### Mass spectrometry

Liquid chromatography tandem mass spectrometry (LC-MS/MS) was performed on a Lumos Tribrid Orbitrap Mass Spectrometer (Thermo Fischer) equipped with Ultimate 3000 (Thermo Fisher) nano-HPLC. Peptides were separated onto a 150-µm inner diameter microcapillary trapping column, packed with approximately two cm of C18 Reprosil resin (5 µm, 100 Å, Dr. Maisch GmbH, Germany), followed by separation on a 50-cm analytical column (PharmaFluidics, Gent, Belgium). Separation was achieved through applying a gradient from 5– 27% ACN in 0.1% formic acid over 90 min at 200 nL min^-1^. Electrospray ionization was enabled through applying a voltage of 2 kV using a home-made electrode junction at the end of the microcapillary column and sprayed from metal tips (PepSep, Denmark). Mass spectrometry survey scan was performed in the Orbitrap, in a range of 400–1,800 m/z at a resolution of 6 x 10^4^, followed by the selection of the 20 most intense ions (TOP20) for CID-MS2 fragmentation in the ion trap using a precursor isolation width window of 2 m/z, AGC setting of 10,000, and a maximum ion accumulation of 100 ms. Singly charged ion species were not subjected to CID fragmentation. Normalized collision energy was set to 35 V and an activation time of 10 ms. Ions in a 10 ppm m/z window around ions selected for MS2 were excluded from further selection for fragmentation for 60 s.

Raw data were analyzed with Proteome Discoverer 2.4 (Thermo Scientific). Assignment of MS/MS spectra was performed using the Sequest HT algorithm by searching the data against a protein sequence database including all entries from our Uniport_Human2018_SPonly database^48^ as well as other known contaminants such as human keratins and common laboratory contaminants. Quantitative analysis between samples was performed by LFQ (label-free quantitation) between different sets of samples. Sequest HT searches were performed using a 10 ppm precursor ion tolerance and requiring each peptides N-/C termini to adhere with Trypsin protease specificity, while allowing up to two missed cleavages. Methionine oxidation (+15.99492 Da) was set as variable modification. A MS2 spectra assignment false discovery rate (FDR) of 1% on both protein and peptide level was achieved by applying the target-decoy database search by user of Percolator.

### Proteomic data preprocessing

Starting with protein level LFQ reports generated from Proteome Discoverer, we performed the following preprocessing steps before inputting the data into Genoppi: (1) performed log_2_ transformation and median normalization of protein intensity values in each experimental sample; (2) filtered out contaminants and protein entries supported by < 2 unique peptides; (3) imputed missing values by randomly sampling from a normal distribution with width of 0.3 standard deviation (SD) and down-shift of 1.8 SD compared to the observed intensity^49^; (4) calculated log_2_ FC for each pair of replicate samples (e.g. bait vs. IgG control). All preprocessed data and subsequent analysis results (average log_2_ FC, *P*-value, and FDR calculated in Genoppi) can be found in **Supplementary Tables 2** and **4**.

### Lists of disease-associated genes

In order to investigate the overlap between interactors identified in our proteomic data and disease-associated genes, we surveyed published literature of genetic studies. For cancer, we used a list of 260 genes identified by exome sequencing^23^ (**Supplementary Table 3**). For neuropsychiatric disease, we aggregated a total of 603 unique genes that have been implicated in autism spectrum disorder (ASD) or schizophrenia (SCZ) genetic studies (**Supplementary Table 3**). This list includes ASD genes identified by exome sequencing^24^ and genes mapped from genome-wide significant loci identified in GWAS of ASD^25^ or SCZ^26^ using Genoppi’s SNP-to-gene mapping framework. For amyotrophic lateral sclerosis (ALS), we curated a list of 53 genes based on literature review^27,28^ (**Supplementary Table 3**).

## Figure Legends

**Supplementary Note 1. Discussion of MDM2 and PTEN IP-MS/MS results.**

To exemplify the analytic features of Genoppi, we generated proteomics data from immunoprecipitations (IPs). We selected four proteins of interest (BCL2, TDP-43, MDM2, PTEN; hereafter called baits), and identified interactomes using label-free liquid chromatography tandem mass spectrometry (LC-MS/MS, Online Methods) in four distinct cell lines (GPiN, G401, T47D, A375; **Fig. 1a**). The presence of the baits in IPs was confirmed by western blot analysis (**Supplementary Fig. 1a**, showing one biological replicate per bait). Although all four baits have been extensively studied^1-4^, their characterization is often limited to a single cell type or disease model. The cell-specific protein-protein interactions of BCL2 and TDP-43 are described in the main text, while here, we focus on the investigated cell-specific interactomes of MDM2 and PTEN. Both proteins are known for their implication in cancer but have not been characterized in neurons^1,2,5-7^.

MDM2 is an important negative regulator of the p53 tumor suppressor, both by inhibiting p53 transcriptional activation and targeting p53 for degradation by the proteasome^1^. More recently, it has been hypothesized that MDM2 is also able to function as an oncogene in p53-independent tumors^5^. In order to investigate this possibility, we immunoprecipitated MDM2 in human epithelial cancer cell lines with a mutated p53 (T47D) and wild-type p53 (A375). The corresponding interactomes were resolved in LC-MS/MS and Genoppi was used to determine the correlation coefficients between IP replicates and the enrichment of MDM2 interactors in each cell line (**Supplementary Fig. 1b**). Proteins interacting with MDM2 were predominantly found in A375, in accordance with the close dependence between p53 and MDM2 in the cell. To identify known protein interactors of MDM2, interactors catalogued in InWeb_InBioMap were overlaid with the experimental results (**Supplementary Fig. 1d**). Interestingly, known interactors of MDM2 are evenly distributed across both cell lines, suggesting that the absence of p53 does not detectably affect the known MDM2 interactome in the studied cancer cell lines. We further employed Genoppi to test for enrichment of cancer genes and found them to be predominantly present among the interactors specific for A375 cells (wild-type p53), although not significantly enriched (**Supplementary Fig. 1e**). This finding indicates that the newly identified interaction partners of MDM2 may depend on p53, a somewhat expected result considering the vast p53-dependent network of oncogenes and the MDM2 regulatory role of p53.

Future research should be aimed at interrogating MDM2 protein networks for newly discovered interactions that can be targeted to interfere with the MDM2-p53 relationship. An immediate testable hypothesis emerging from this analysis is to evaluate whether any of the MDM2 interactors shown here can be targeted to lower the existing MDM2-p53 connection, and used to treat cancers. Conversely, thorough characterization of T47D specific interactors is key to decipher the p53-independent protein network of MDM2.

PTEN (a well-known tumor suppressor gene that antagonizes phosphatidylinositol 3-phosphate kinase [PI3K] and AKT signaling) encodes for a phosphatase and is mutated in a large number of cancers at a high frequency^2^. Interestingly, germline mutations of PTEN have also been described in a subset of patients with autism spectrum disorders and macrocephaly^6^. However, it is unclear whether the role of PTEN in cell proliferation is also implicated in these neurodevelopmental phenotypes^6,7^. To investigate this possibility, we compared the interactomes of PTEN in human cancer cells (G401) and neurons (iPSC-derived glutamatergic patterned induced neurons, GPiNs). IP replicates comparing the two cell lines correlate highly (≥ 0.69; **Supplementary Fig. 1c**) and more known PTEN interactors from InWeb_InBioMap were identified in G401 cells (**Supplementary Fig. 1f**), suggesting that PTEN does not interact with its classic partners in human neurons. Next, we used Genoppi to test the representation of cancer genes in both cell lines; in line with the expectation that PTEN interaction partners are linked to proliferation in both cancer cells and human neurons, we did not observe a significant difference between cell lines (**Supplementary Table 3** and **Supplementary Fig. 1g**). Genoppi was also used to assess the enrichment of proteins encoded by risk genes previously linked to neuropsychiatric disease including autism and schizophrenia (**Supplementary Table 3** and **Supplementary Fig. 1h**). We found a comparable distribution of risk gene products in G401 cells and GPiNs, indicating that they are unlikely part of pathways that drive the neuron-specific function of PTEN. This result highlights the importance of studying and comparing both cell-type-specific and shared interactions of PTEN in cancer cells and human neurons to yield insights into its specific function in neurodevelopment and disease.

---

**Supplementary Figure 1.**
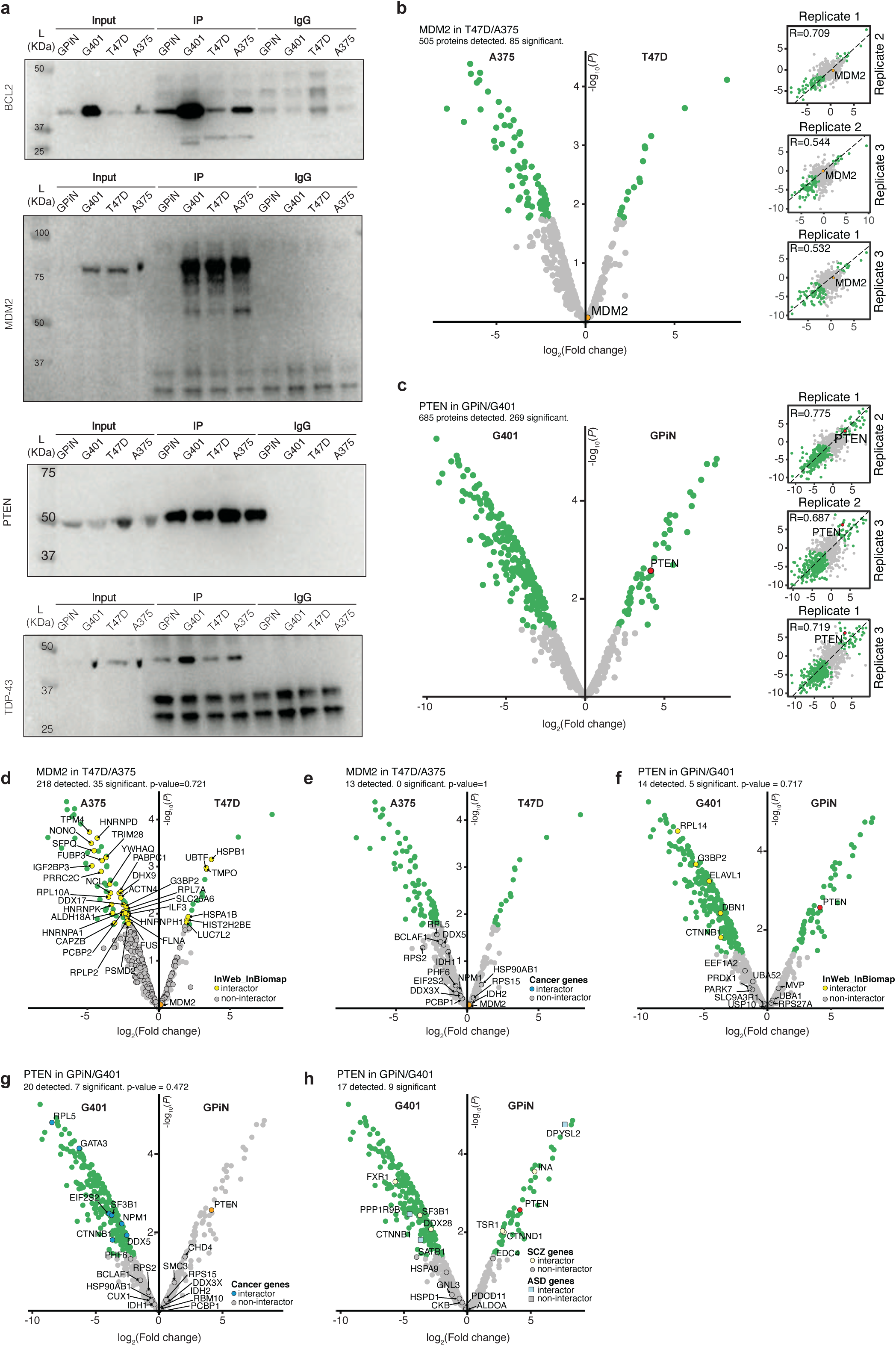
Additional IP-MS/MS analysis results (related to **Figure 2**). **(a)** Western blots show IP results for all baits in all cell lines; IN, FT, IP, and IgG stand for input, flow-through, immunoprecipitation, and IgG isotype control, respectively. **(b)** MDM2 IP results in T47D versus A375 cells. Left: volcano plot showing average log_2_ fold change (FC) and corresponding P-value of each identified protein. Right: scatter plots showing reproducibility of three IP replicates in terms of log_2_ FC correlation; Pearson correlation (r) is reported in each plot. The bait (MDM2), interactors (log_2_ FC > 0 and FDR ≤ 0.1), and non-interactors are shown in orange, green, and grey, respectively. **(c)** PTEN IP results in GPiN versus G401 cells. Same layout as in (b), except the bait (PTEN) is shown in red. **(d)-(e)** Volcano plot from (b) overlaid with known MDM2 interactors in InWeb_InBioMap (d) or cancer genes (e); overlaid proteins are marked by black border circles. **(f)-(h)** Volcano plot from (c) overlaid with known PTEN interactors in InWeb_InBioMap (f), cancer genes (g), or neuropsychiatric disease genes (h); overlaid proteins are marked by black border circles or squares.

**Supplementary Figure 2.**
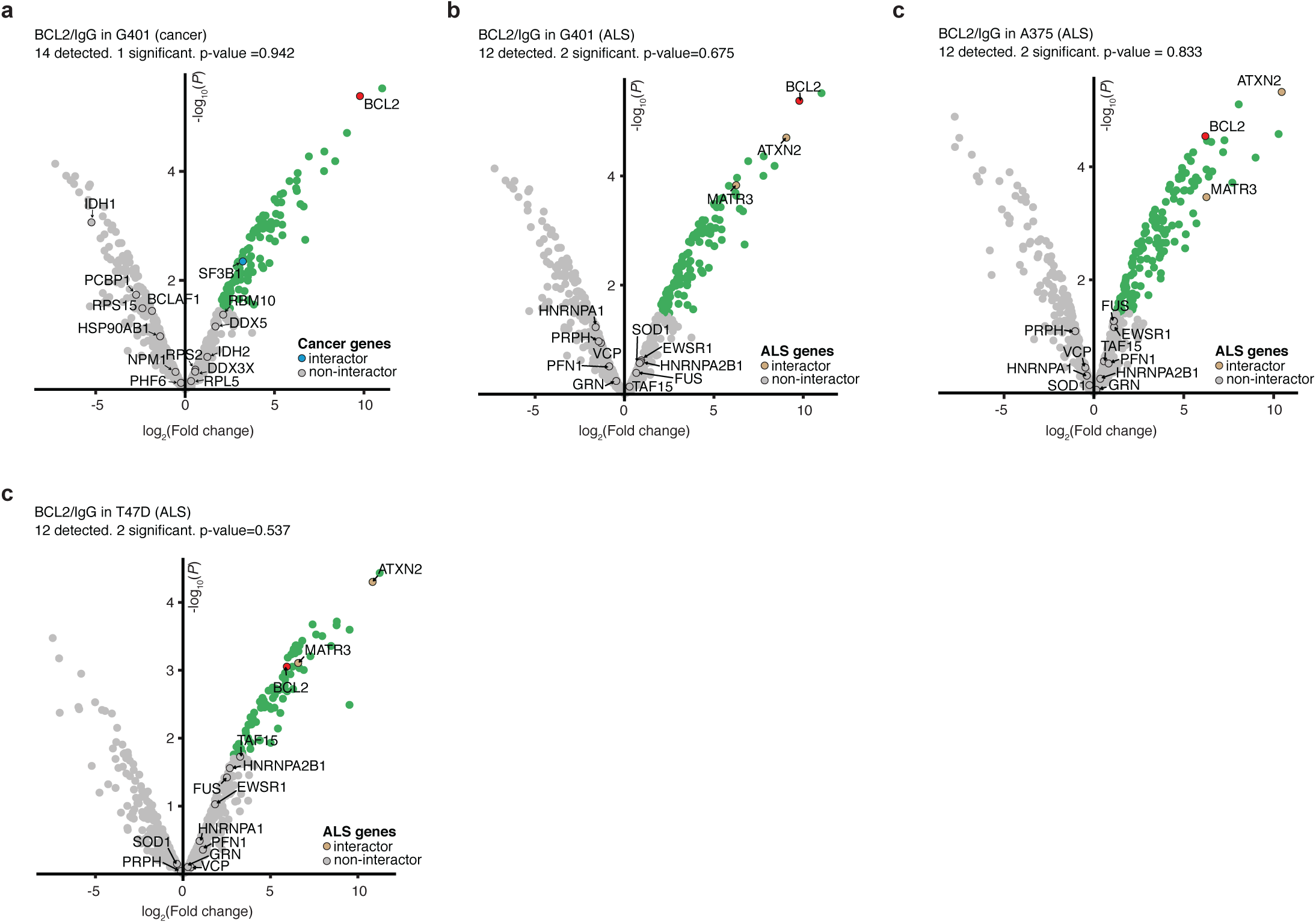
BCL2 versus IgG control IP results in G401 **(a-b)**, A375 **(c)**, and T47D **(d)** cells. The volcano plots show average log_2_ fold change (FC) and corresponding -log_10_ P-value of each identified protein; the bait (BCL2), interactors (log_2_ FC > 0 and FDR ≤ 0.1), and non-interactors are shown in red, green, and grey, respectively. Each volcano plot is overlaid with proteins encoded by either cancer genes **(a)** or ALS genes **(b-d)**, with corresponding overlap enrichment P-value reported in each panel; overlaid proteins are marked by black border circles.

**Supplementary Figure 3.**
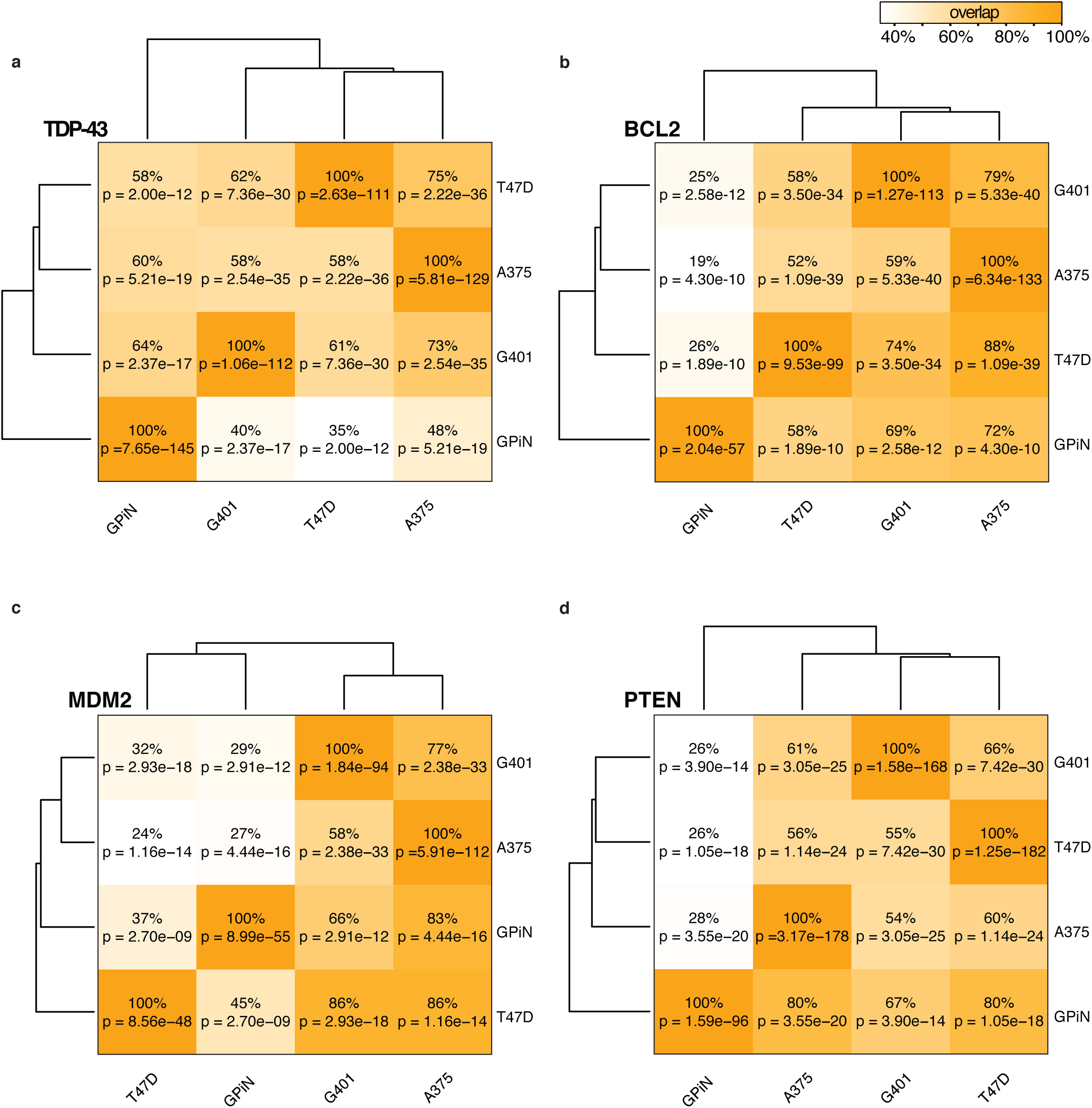
Overlap of interactors identified for each bait between each pair of cell lines. Clustered heatmaps indicating the percentage of overlap between significant interactors (log_2_ FC > 0 and FDR ≤ 0.1) of baits TDP-43 **(a)**, BCL2 **(b)**, MDM2 **(c)**, and PTEN **(d)** when compared against IgG control across different cell types. The overlap percentages were calculated using the row cell type as reference; the corresponding P-values were calculated using a one-tailed hypergeometric test. Dendrograms were generated from hierarchical clustering of the overlap percentages.

## Supplementary Protocol

### Getting started

Genoppi is an open-source software for performing quality control and analyzing quantitative proteomic data. Genoppi streamlines the integration of proteomic data with external datasets such as known protein-protein interactions in published literature, data from genetic studies, gene set annotations, or other user-defined inputs. This protocol provides documentation for using the interactive Genoppi web application. Source code for the application and the stand-alone Genoppi R package can be downloaded at github.com/lagelab/Genoppi for local installation.

Please direct questions and comments to Kasper Lage.

### Input format

Genoppi can be used to analyze quantitative proteomic data that contain protein enrichment values between studied conditions, such as bait vs. control immunoprecipitations followed by mass spectrometry (IP-MS). Protein quantification results generated using labeling-based (e.g., iTRAQ, TMT, or SILAC) or label-free MS methods can be inputted into Genoppi following the input file format described below.

The input file must be a tab-delimited text file. At minimum, the file must contain three columns, with one column specifying protein identifiers and two columns listing protein enrichment values for two or more experimental replicates. More specifically:

#### Column 1

protein identifiers as either HUGO Gene Nomenclature Committee^1^ (HGNC) [www.genenames.org] approved symbols (with “gene” as column header), or UniProt^2^ [www.uniprot.org] accession numbers (with “accession_number” as column header).

#### Columns 2, 3, (+ optional additional columns)

log_2_ fold change (FC) values for ≥ two replicates, with “rep1”, “rep2”, and so on as column headers for the replicates.

**OR**

#### Columns 2, 3, 4

average log_2_ FC across replicates (“logFC”) with corresponding *P*-value (“pvalue”) and false discovery rate (“FDR”) calculated using a statistical test (e.g. a moderated t-test).

Missing values are not allowed; any rows with missing values would be disregarded with no error message.

Examples of accepted input format with correct column headers:

**1. HGNC symbol and log**_**2**_ **FC for two replicates**

| gene | rep1 | rep2 |
| --- | --- | --- |
| FOXP2 | -0.496 | -0.546 |
| RB1 | 0.402 | 0.265 |
| SHH | 0.08 | 0.104 |

**2. UniProt accession number and log**_**2**_ **FC for three replicates**

| accession_number | rep1 | rep2 | rep3 |
| --- | --- | --- | --- |
| O15409 | -0.496 | -0.546 | -0.447 |
| P06400 | 0.402 | 0.265 | 0.410 |
| Q15465 | 0.08 | 0.104 | 0.125 |

**3. HGNC symbol and pre-calculated results of statistical test**

| gene | logFC | pvalue | FDR |
| --- | --- | --- | --- |
| FOXP2 | -0.521 | 1.64e-3 | 4.14e-3 |
| RB1 | 0.334 | 0.0118 | 0.0189 |
| SHH | 0.092 | 0.211 | 0.242 |

For Mac users exporting data from Excel format, please convert it to text file by selecting “File” > “Save As…” > “File Format” > “Tab delimited Text (.txt)”. This would avoid generating a file that terminates each line with a carriage return character, which is incompatible with subsequent analysis in Genoppi.

### Basic plotting

**Screenshot 1** illustrates the basic user interface of the Genoppi application. After the user uploads a “Single File” input in the left panel (**Screenshot 1a**), the “Basic plotting” module will generate an interactive volcano plot, depicting the average log_2_ FC of proteins on the x-axis and the -log_10_ *P*-value of enrichment on the y-axis. If log_2_ FC values from ≥ two replicates are provided in the input file, a moderated *t-*test from the limma^3^ R package is applied to calculate the average log_2_ FC, nominal *P*-value, and FDR (see **Online Methods**); otherwise, Genoppi uses the user-supplied statistics to generate the plot. In addition, a scatter plot showing replicate log_2_ FC correlation is generated if the input file includes separate replicates; when there are > 2 replicates, the user can select from a drop-down menu to show the scatter plot corresponding to each pair of replicates (**Screenshot 1b**).

In the default coloring scheme, enriched proteins with log_2_ FC ≥ 0 and FDR ≤ 0.1 are in green, and other detected proteins are in grey. The user can change the colors (**Screenshot 1c**) or modify the significance threshold for defining enriched proteins based on different FDR, *P*-value, and log_2_ FC cutoffs (**Screenshot 1d**). The “Summary” box shows the number of enriched proteins (and total number of detected proteins) based on the specified threshold, as well as the correlation between replicates when appropriate. Hovering over each protein’s data point in either the volcano or scatter plot would show its corresponding HGNC symbol. The user can also query specific HGNC symbols to label bait and other proteins in the plots (**Screenshot 1e**).

**Screenshot 1.**
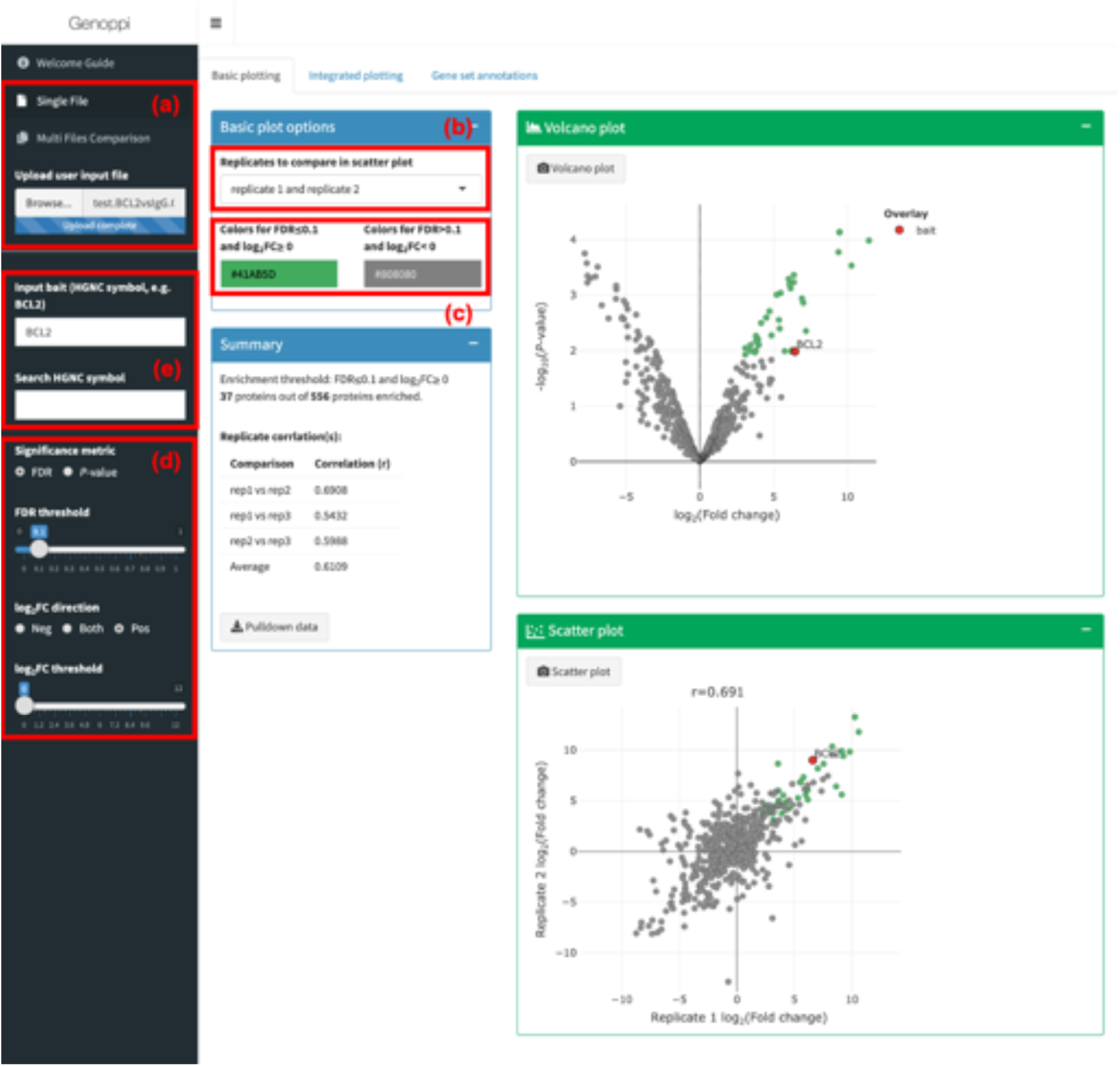
Basic plotting interface showing volcano and replicate correlation scatter plots generated from input proteomic data.

### Integrated plotting

In the “Integrated plotting” module, Genoppi enables integration of the proteomic data with data from the InWeb_InBioMap database^4^, NHGRI-EBI GWAS catalog^5^, gnomAD database^6^, or user-uploaded SNP or gene lists.

#### InWeb_InBioMap

Genoppi can overlay the proteomic data with data from InWeb_InBioMap (InWeb) [www.intomics.com/inbio/map.html#downloads], which contains human protein-protein interactions compiled from > 40,000 published articles. This integration enables the user to easily distinguish new interactions from those already reported in the literature. The user can search for InWeb interactors of a specific protein to visualize their overlap with enriched proteins in the proteomic data in an overlaid volcano plot (**Screenshot 2a**). In this plot, InWeb interactors (as well as other data types described below) detected in the proteomic data are labeled using an adjustable color and shape scheme (**Screenshot 2b**).

#### GWAS catalog

Genoppi can perform SNP-to-gene mapping for SNPs found in the 1000 Genomes Project^7^ [www.internationalgenome.org], using pre-calculated pairwise linkage disequilibrium (LD) measures between SNPs to identify all genes in LD regions (see **Online Methods**). Therefore, the user can query diseases and traits found in the NHGRI-EBI GWAS catalog [www.ebi.ac.uk/gwas] to identify genes mapped from published trait-associated SNPs in the catalog (**Screenshot 2c**). Proteins encoded by the mapped genes would be labeled in the interactive volcano plot, and hovering over each of these proteins would show the SNP(s) that map to it.

#### gnomAD

Genoppi can also identify proteins encoded by genes that are likely intolerant of loss-of-function (LoF) mutations using constraint data from gnomAD [gnomad.broadinstitute.org]. The user can label proteins with pLI scores (i.e. probability of intolerance to LoF mutations) greater than an adjustable threshold to visualize the most intolerant proteins in the overlaid volcano plot (**Screenshot 2d**).

**Screenshot 2.**
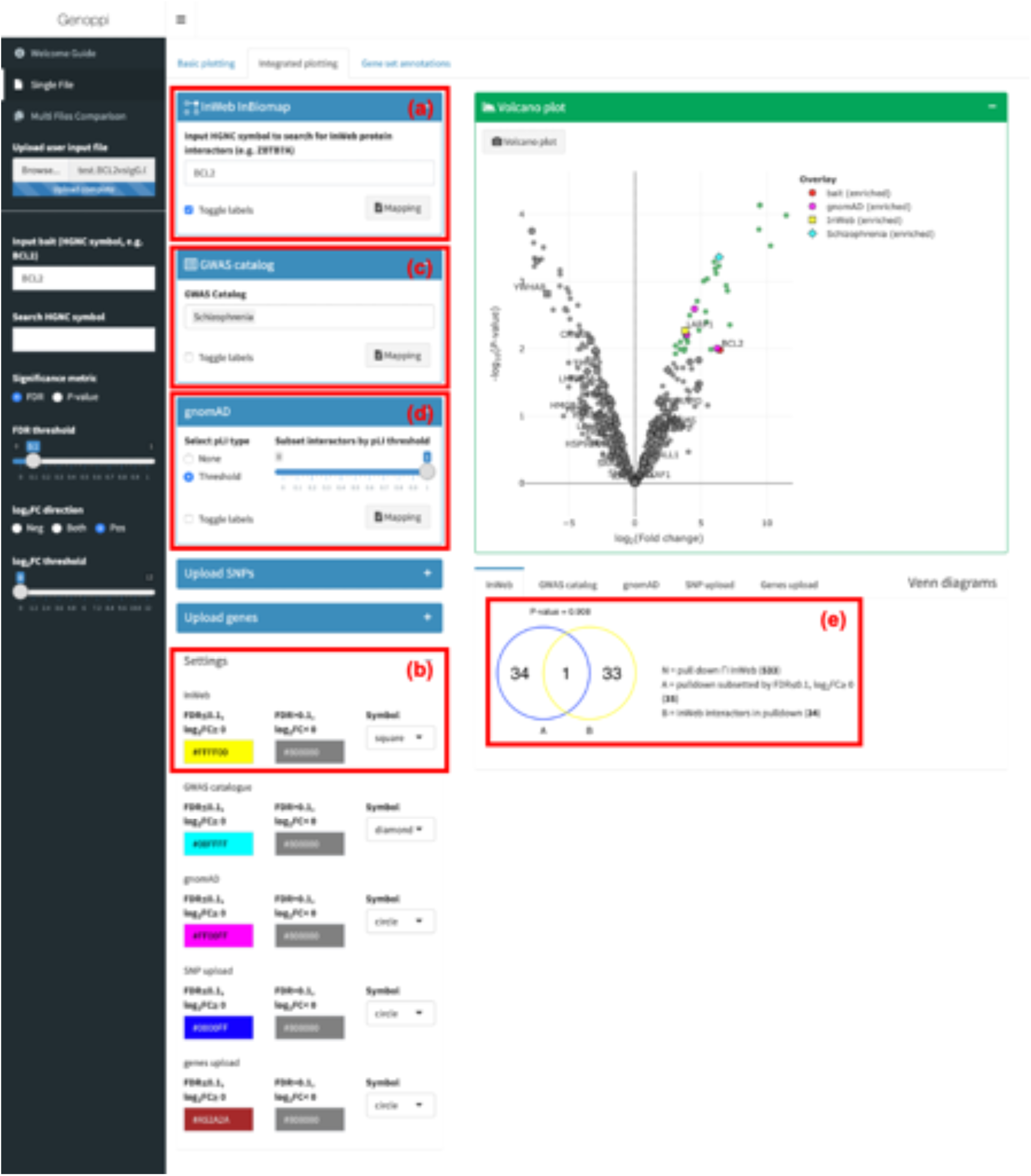
Integrated plotting interface showing integration of proteomic data with InWeb, GWAS catalog, and gnomAD data.

#### Upload SNPs or genes

Besides incorporating the public datasets described above, the user may also upload one or more custom SNP or gene lists (e.g. disease-causing genes curated from literature review or genes implicated by gene-based burden testing) to assess their overlaps with the proteomic data (**Screenshot 3a**). As described in the “GWAS catalog” section, uploaded SNPs would be mapped to genes in LD using Genoppi’s built-in SNP-to-gene mapping functionality. The SNP list(s) must be uploaded as a tab-delimited plain text file containing two columns: “listName” (name for each list) and “SNP” (rsID). For example:

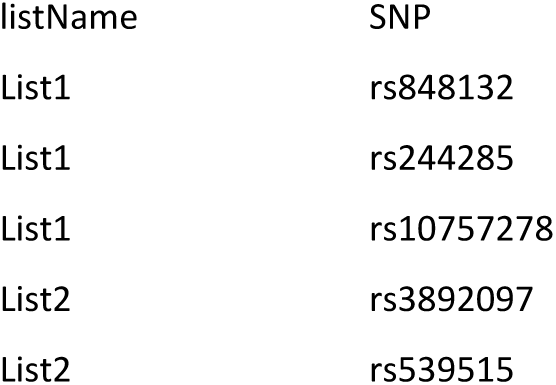

Similarly, the gene list(s) must be uploaded as a tab-delimited text file consisting of two columns: “listName” (name for each list) and “gene” (HGNC symbol). For example:

| listName | gene |
| --- | --- |
| ListA | SHH |
| ListA | UBC |
| ListB | FOXP2 |
| ListB | RB1 |
| ListB | KRAS |

In the “Integrated plotting” module, the user may input any combination of InWeb interaction partners, GWAS catalog mapped genes, gnomAD constrained genes, and SNP and gene lists. The resulting volcano plot would highlight all identified proteins from these inputs. Overlaying multiple datasets could result in a densely labeled plot, in which case the user can choose to remove text labels for the proteins using the “Toggle labels” option for each data type (**Screenshot 3b**).

**Screenshot 3.**
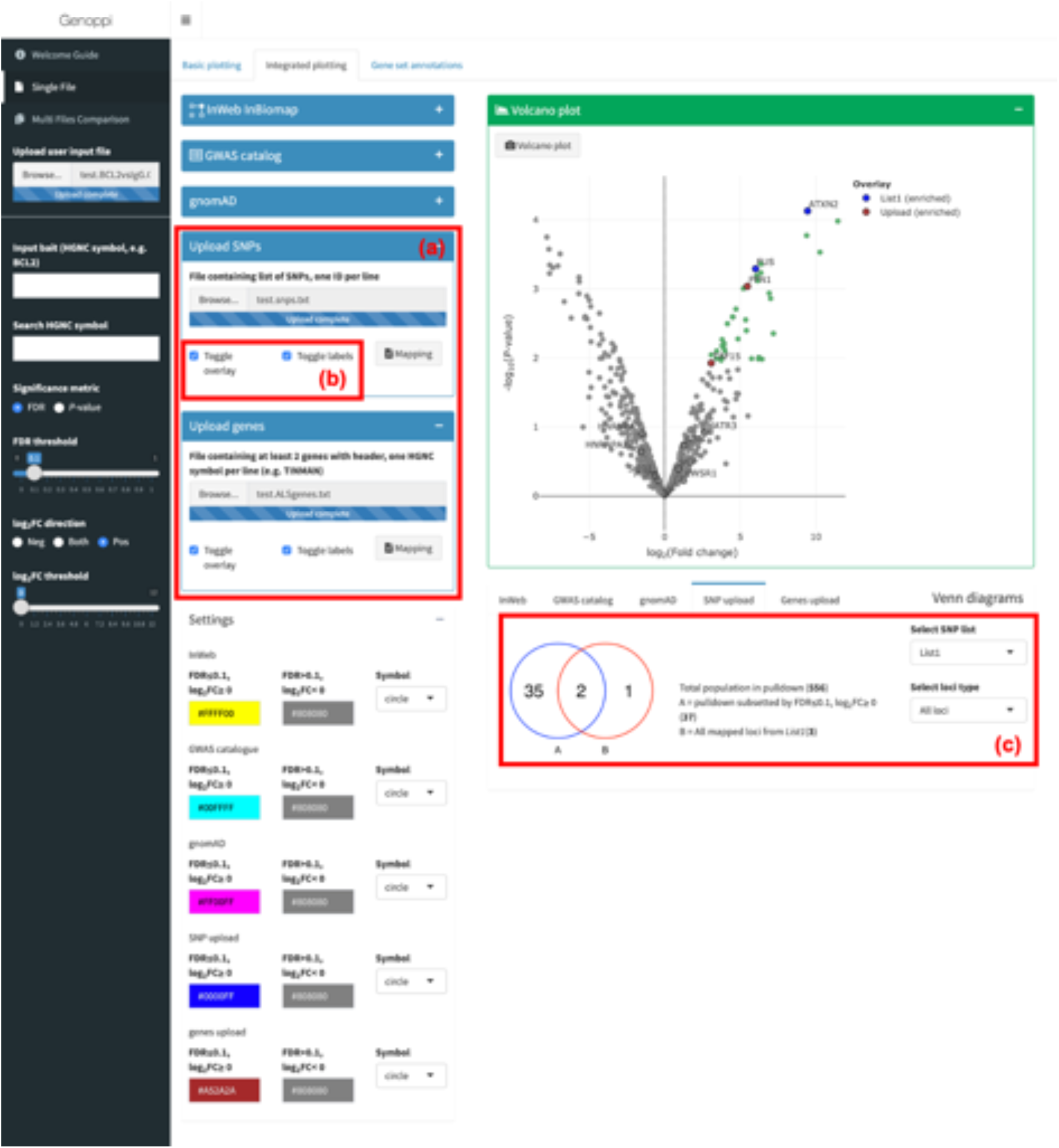
Integrated plotting interface showing integration of proteomic data with user-uploaded SNP and gene lists.

#### Venn diagrams

In the “Integrated plotting” module, Genoppi also summarizes the overlaps between the enriched proteins in the proteomic data and the various data types described above using Venn diagrams. When showing overlap with InWeb_InBioMap, gnomAD constrained genes, or user-uploaded gene lists, Genoppi also assesses the overlap enrichment by calculating a hypergeometric *P*-value, which is displayed above the Venn diagram (**Screenshot 2e**; see **Online Methods**). For genes mapped from GWAS catalog or user-uploaded SNP lists, this calculation is not performed as the statistic is not robust when each SNP could be mapped to multiple genes in LD. In addition, for user-uploaded SNP lists (which should contain only independent SNPs), Genoppi generates three Venn diagrams to show the overlap of enriched proteins with all mapped genes, genes in single-gene loci, or genes in multi-gene loci, respectively; these diagrams can be selected using a drop-down menu (**Screenshot 3c**).

If the user uploads multiple SNP or gene lists, Venn diagrams and overlap statistics for each list would be generated separately for each list, and the individual list results can be accessed through clicking on the list name in a drop-down menu (**Screenshot 3c**). Note that when a bait protein has been indicated in the “Input bait” search box in the left panel, Genoppi would exclude the bait when calculating the numbers and statistics in the *Venn diagrams* section.

### Gene set annotations

In the “Gene set annotations” module, Genoppi enables annotation of the proteomic data with gene sets from various databases, including HGNC gene groups, Gene Ontology^8,9^ (GO) [geneontology.org] terms (molecular function, cellular component, and biological process), and MSigDB^10,11^ [www.gsea-msigdb.org/gsea/msigdb/index.jsp] gene sets (H and C1-C7 collections), allowing the user to explore the diversity of protein functions in the proteomic results.

The user can annotate enriched proteins in their volcano plot by selecting a collection of gene sets from a drop-down menu (**Screenshot 4a**). Proteins belonging to different gene sets are annotated using square markers of distinct colors; the marker size is scaled with the frequency of each gene set (i.e. number of proteins assigned to each set), providing quick visualization of overrepresentation trends in the data. The volcano plot can display up to 100 most recurrent gene sets at once; the user can further filter these top gene sets using the frequency slider (**Screenshot 4a**). Hovering over each marker in the resulting volcano plot would show the protein’s gene set annotations. Alternatively, the table below the volcano plot lists all the gene set annotations without the 100 gene sets limitation. Finally, the user can also query specific gene sets using a search box (**Screenshot 4b**), and the proteins belonging to the queried sets would be labeled with diamond markers in the volcano plot.

**Screenshot 4.**
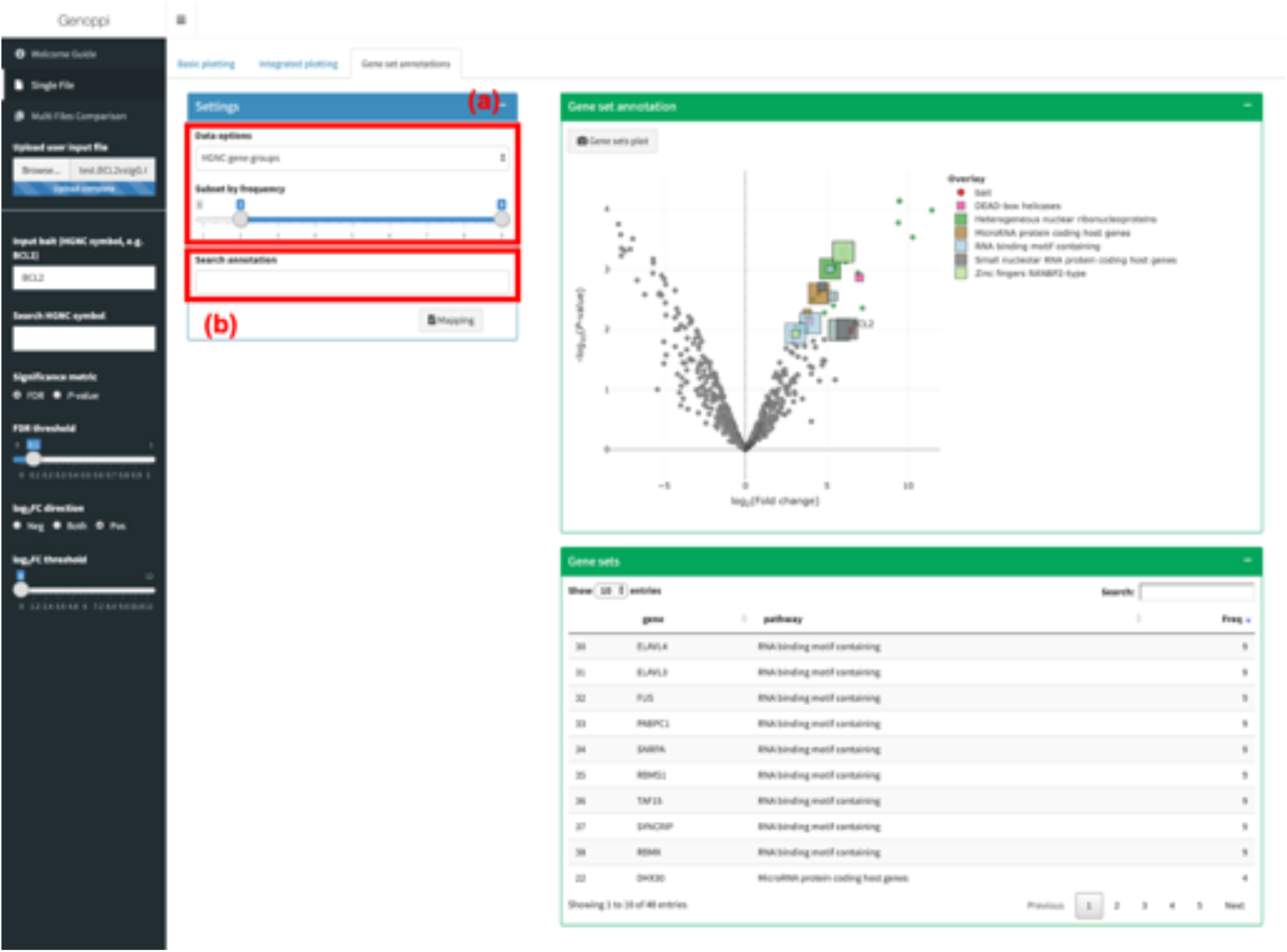
Gene set annotations interface showing enriched proteins in proteomic data annotated with most recurrent HGNC gene groups.

### Multiple files comparison

Besides performing analyses for a single proteomic dataset as described in the previous sections, Genoppi also allows comparison of multiple proteomic datasets at once. Using the “Multi Files Comparison” input option, the user can upload two to three proteomic datasets to perform comparative analyses. In the “Basic plotting”, “Integrated plotting”, and “Gene set annotations” modules, Genoppi would generate side-by-side volcano, scatter, and/or Venn diagram plots to compare the multiple datasets and offer customization features similar to that for the “Single File” input.

#### Protein comparison

In the “Protein comparison” module, the user can visualize the overlaps between enriched proteins identified in different datasets. The significance threshold for defining enriched proteins can be individually adjusted for each dataset before generating the comparison results. In the resulting volcano and scatter plots, each enriched protein is color-coded based on the combination of datasets that share this enriched protein. The number of proteins in each combination group is also summarized in a Venn diagram and the identities (i.e. HGNC symbols) of these proteins are displayed in a table.

### Downloads

Individual plots and data files generated by Genoppi can be downloaded in their respective modules by clicking on the interactive download buttons. In general, plots are saved as PNG image files, while text files are saved in comma-separated CSV format. Download buttons are only active once the relevant data have been generated.

